# Clonal transcriptomics identifies mechanisms of chemoresistance and empowers rational design of combination therapies

**DOI:** 10.1101/2021.12.09.471927

**Authors:** Sophia A Wild, Ian G Cannell, Katarzyna Kania, Ashley Nicholls, Dario Bressan, CRUK IMAXT Grand Challenge Team, Gregory J Hannon, Kirsty Sawicka

## Abstract

Tumor heterogeneity is thought to be a major barrier to successful cancer treatment due to the presence of drug resistant clonal lineages. However, identifying the characteristics of such lineages that underpin resistance to therapy has remained challenging. Here we utilize clonal transcriptomics with WILD-seq; **W**holistic **I**nterrogation of **L**ineage **D**ynamics by **seq**uencing, in mouse models of triple-negative breast cancer (TNBC) to understand response and resistance to therapy, including BET bromodomain inhibition and taxane-based chemotherapy. This analysis revealed oxidative stress protection by NRF2 as a major mechanism of taxane resistance and led to the discovery that our tumor models are collaterally sensitive to asparagine deprivation therapy using the clinical stage drug L-asparaginase after frontline treatment with docetaxel. In summary, clonal transcriptomics with WILD-seq identifies mechanisms of resistance to chemotherapy that are also operative in patients and pin points asparagine bioavailability as a druggable vulnerability of taxane resistant lineages.

## Introduction

Intra-tumoral heterogeneity (ITH) is thought to underlie tumor progression and resistance to therapy by providing a reservoir of phenotypically diverse clonal lineages on which selective pressures from the microenvironment or therapeutic intervention exert their effects (Bhang et al., 2015; Turajlic & Swanton, 2016). Inference of clonal composition from bulk sequencing has elucidated the breadth of ITH across tumor types and suggests that often rare pre-existing clones can resist therapy-induced killing to drive relapse (Dentro et al., 2021; Ding et al., 2012; Gerlinger et al., 2012; Jamal-Hanjani et al., 2014; Landau et al., 2013). However, such methods are limited by their inability to characterize such resistant clones beyond genotype and how their properties change over time and in response to therapy. Recently, several lineage tracing approaches have emerged that are able to link clonal identity with gene expression by utilizing expressed genetic barcodes that are read-out by single cell RNA sequencing (Biddy et al., 2018; Gutierrez et al., 2021; Quinn et al., 2021; Simeonov et al., 2021; Weinreb et al., 2020; Yang et al., 2022). These powerful methods allow deconvolution of complex mixtures of clones while simultaneously providing a gene expression profile of those cells that can indicate the pathways on which they depend. However, to date in solid tumors these technologies have mostly been used to study drug response *in vitro* (Gutierrez et al., 2021; Oren et al., 2021) or metastatic dissemination *in vivo* (Quinn et al., 2021; Simeonov et al., 2021; Yang et al., 2022) and have not been utilized to study therapeutic response in immune-competent models.

A thorough understanding of the biomarkers of sensitivity and mechanisms of resistance to chemotherapy is essential if we are to improve patient outcomes. Most existing combination cancer therapies are not rationally designed but were instead empirically optimized to avoid overlapping toxicities. More recently alternative therapeutic strategies have emerged including synthetic lethality, drug synergy (Al-Lazikani et al., 2012; O’Neil et al., 2017) and collateral sensitivity (Mueller et al., 2021; Pluchino et al., 2012; Zhao et al., 2016) that aim to leverage selective vulnerabilities of tumor cells while minimizing toxicity. Of particular promise is collateral sensitivity, in which as a tumor becomes resistant to one drug it comes at the cost of sensitivity to a second drug. Since many modern clinical trials occur in the context of neo-adjuvant chemotherapy, the identification of frontline therapy-induced collateral sensitivities to second line therapy would have the potential to be rapidly translated into improved outcomes for patients.

Here we develop WILD-seq (**W**holistic **I**nterrogation of **L**ineage **D**ynamics by **seq**uencing), an accessible and adaptable platform for lineage tracing at the single-cell transcriptomic level that facilitates *in vivo* analysis of clonal dynamics and apply it to the study of syngeneic triple negative breast cancer (TNBC) mouse models. Our optimized pipeline ensures recurrent representation of clonal lineages across animals and samples, facilitating analysis of clonal dynamics under the selective pressure of therapeutic intervention. Importantly, analysis of response of TNBC models to frontline taxane-based chemotherapy revealed an enrichment of clones with high levels of NRF2 signaling, implicating defense against oxidative damage as a major determinant of resistance to chemotherapy. Building on the work of others (LeBoeuf et al., 2020) we show that these NRF2-high, taxane-resistant, lineages are collaterally sensitive to asparagine deprivation with L-asparaginase and that they adapt to this second line intervention by up-regulating *de novo* asparagine synthesis through asparagine synthetase (Asns). Together these data indicate that high levels of NRF2 signaling, which is also observed in patients following neo-adjuvant chemotherapy, promotes both resistance to chemotherapy and sensitivity to asparagine deprivation and warrant the exploration of L-asparaginase as a therapeutic modality in solid tumors.

## Results

### Establishment of an expressed barcode system to simultaneously detect clonal lineage and gene expression

WILD-seq uses a lentiviral library to label cells with an expressed, heritable barcode that enables identification of clonal lineage in conjunction with single cell RNA sequencing. The WILD-seq construct comprises a zsGreen transcript which harbours in its 3’ untranslated region (UTR) a barcode consisting of two 12 nucleotide variable regions separated by a constant linker (Fig. 1a). Each variable region is separated from any other sequence in the library by a Hamming distance of 5 to allow for library preparation and sequencing error correction and our library contains over 1.5 million unique barcodes. The barcode is appropriately positioned relative to the polyadenylation signal to ensure its capture and sequencing by standard oligo-dT single cell sequencing platforms.

**Figure 1.**
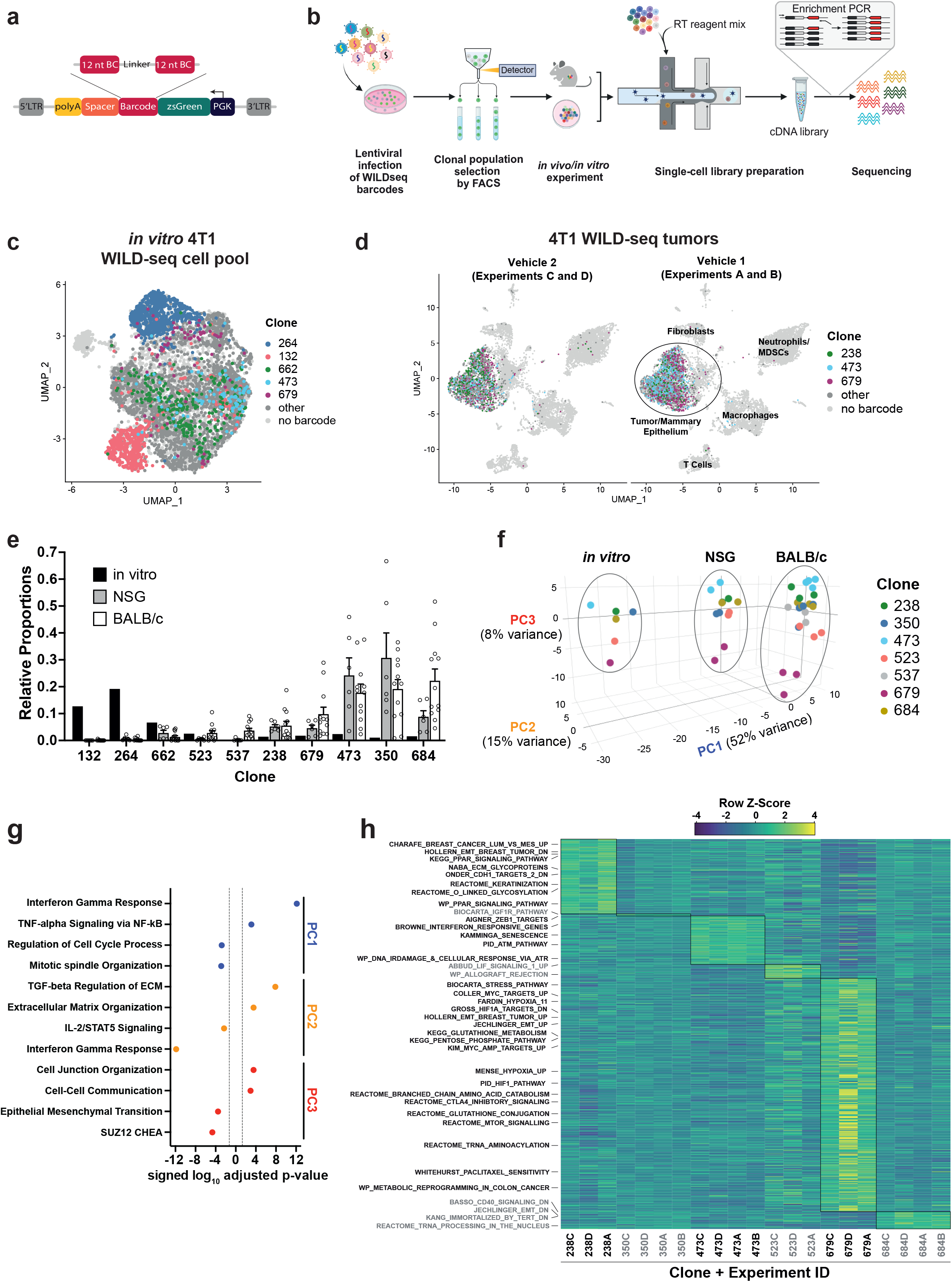
Establishment of an expressed barcode system to simultaneously detect clonal lineage and gene expression from single cells *in vivo*. **a. Lentiviral construct design.** A PGK promoter drives expression of a transcript encoding zsGreen harboring a WILD-seq barcode sequence in the 3’UTR. A spacer sequence and polyadenylation signal ensure that that the barcode is detectable as part of a standard oligo dT single cell RNA library preparation and sequencing pipeline. The barcode cassette comprises 2 distinct 12 nucleotide barcode sequences separated by a constant 20 nucleotide linker region. The library of barcode sequences was designed with Hamming distance 5 to allow for sequencing error correction. **b. Schematic of WILD-seq method.** Tumor cells are infected with the WILD-seq lentiviral library and an appropriate size population of zsGreen positive cells isolated, each of which will express a single unique WILD-seq barcode. This WILD-seq barcoded, heterogenous cell pool is then subjected to an intervention of interest (such as *in vivo* treatment of the implanted pool with a therapeutic agent) and subsequently analyzed by single cell RNA sequencing using the 10X Genomics platform. An additional PCR amplification step is included that specifically enriches for the barcode sequence to increase the number of cells to which a WILD-seq barcode can be conclusively assigned. **c. scRNA-seq of *in vitro* 4T1 WILD-seq cell pool.** UMAP plot of *in vitro* cultured 4T1 WILD-seq cells. Cells for which a WILD-seq clonal barcode is identified are shown as dark grey or colored spots. Cells which belong to five selected clonal lineages are highlighted. **d. scRNA-seq of 4T1 WILD-seq tumors.** UMAP plots of vehicle-treated 4T1 WILD-seq tumors generated by injecting the 4T1 WILD-seq pool into the mammary fatpad of BALB/c mice. Four independent experiments were performed each involving injection into 3 separate host animals. Six animals from experiments A and B received vehicle 1 (10% DMSO, 0.9%β-cyclodextrin) and six animals from experiments C and D received vehicle 2 (12.5% ethanol, 12.5% Kolliphor). **e. Clonal representation.** Proportion of tumor cells assigned to each clonal lineage based on the WILD-seq barcode (n = 1 for *in vitro* cultured cells, n = 6 for tumors from NSG mice, n = 12 for vehicle-treated tumors from BALB/c mice). Selected clones from the most abundant lineages are plotted. Data represents mean ± SEM. **f. Principal component analysis of clonal transcriptomes.** Pseudo-bulk analysis was performed by summing counts for all tumor cells expressing the same WILD-seq clonal barcode within an independent experiment. For *in vivo* tumor samples each point represents the combined cells from 3 animals. Principal component analysis of normalized pseudo-bulk count data showed separation of samples by origin with PC1 and PC2 and separation by clonality with PC3. **g. Transcriptomic programs associated with principal components.** The top/bottom 50 gene loadings of PC1, PC2 and PC3 were analyzed using Enrichr (Chen et al., 2013; Kuleshov et al., 2016; Xie et al., 2021). **h. Clonal transcriptomic signatures from vehicle-treated BALB/c tumors.** An AUCell score (Aibar et al., 2017) enrichment was calculated for each clone and for each experiment by comparing cells of a specific clonal lineage of interest to all assigned tumor cells within the same experiment. All gene sets which showed consistent and statistically significant enrichment in one of the six most abundant clones across experiments are illustrated.

The standard WILD-seq pipeline is illustrated in Figure 1b. A heterogeneous cell line is transduced with a barcode library at low multiplicity of infection (MOI) to ensure that each cell receives a maximum of one barcode. An appropriate size pool of barcoded clones is selected and stabilized in culture. Empirically, we have found a pool established from 750 individual clones works well to provide effective representation of the diversity within the cell lines used herein while also enabling recurrent representation of the same clones across animals and experiments. Once stabilized in culture, the pool of WILD-seq clones can be analyzed directly by single cell sequencing or injected into a recipient animal for *in vivo* tumor growth. WILD-seq single cell sequencing libraries can be prepared using a standard oligo-dT based protocol and addition of an extra PCR amplification step can be used to increase coverage of the barcode region and aid cell lineage assignment.

We first established a WILD-seq clonal pool from the mouse 4T1 cell line, a triple negative mammary carcinoma model that can be orthotopically implanted into the mammary fat pad of a BALB/c syngeneic host, which we have previously shown to be heterogeneous with distinct sub-clones having unique biological properties (Wagenblast et al., 2015). We performed single cell sequencing of the *in vitro* WILD-seq pool (Fig. 1c) and *in vivo* tumors derived from this clonal pool (Fig. 1d). Over the course of our studies, we injected multiple cohorts of mice with our WILD-seq 4T1 pool as detailed in Supplementary Table 1, some of which were subjected to a specific drug regime. All tumors were harvested at humane endpoint, as determined by tumor volume unless otherwise stated and immediately dissociated for single cell sequencing.

For the purpose of characterising the baseline properties of our clones, we performed an in-depth transcriptomic analysis of all tumors from untreated and vehicle-treated animals. A WILD-seq barcode and thereby clonal lineage could be unambiguously assigned to 30-60% of cells per sample within the presumptive tumor cell/mammary epithelial cell cluster. 132 different WILD-seq barcodes were observed *in vitro* and in total 94 different WILD-seq barcodes were observed across our *in vivo* tumor samples. Our *in vivo* tumor samples comprised both tumor cells and host cells of the tumor microenvironment including cells of the innate and adaptive immune system, enabling simultaneous profiling of the tumor and its microenvironment (Fig. 1d). Clustering was performed after removal of reads mapping to the WILD-seq vector, to avoid any influence of the WILD-seq transcript on clustering, and the WILD-seq barcode assignment subsequently overlaid onto these data. The tumor cell clusters were clearly identifiable by the high expression of the barcode transcript. Occasionally a barcode was observed in cells which clustered according to their transcriptome outside of the main tumor cluster. Since this could be the result of sequencing or technical error causing a mismatch between the WILD-seq barcode and the cell of origin, only barcoded cells that clustered within the main tumor/mammary epithelium cell cluster were included in our analysis.

We reproducibly observed the same clonal populations across animals and independent experiments which is critical to our ability to examine the effects of different interventions and treatments (Fig. 1d, 1e). The relative abundance of clones was similar in tumors grown in NOD scid gamma (NSG) immunodeficient and BALB/c immunocompetent mice but was drastically different to that found in the *in vitro* cell pool from which they were established (Fig. 1e, Supplementary Table 2), suggesting that clones that show greatest fitness in cell culture do not necessarily show fitness *in vivo*. Therefore, *in vitro* clonal lineage tracking experiments are likely to capture a different collection of clones and have the potential to identify sensitive or resistance clones that are not represented *in vivo*. Pseudo-bulk analysis of the major clonal lineages revealed that the composition of the tumor microenvironment has a dramatic effect on the transcriptome of the tumor cells for all clones (Fig. 1f). Comparison of *in vitro* culture, tumors from NSG mice, and tumors from BALB/c mice by principal component analysis (PCA), showed clear separation of the tumor cells depending on their environment, with differences in interferon gamma signaling, TNF-alpha signaling, and cell cycle being most prominent between cells grown *in vivo* and *in vitro* (PC1, Fig. 1g). Differences in gene expression between tumors growing in immunocompetent and immunodeficient hosts were related to changes in the expression of extracellular matrix proteins and changes in interferon gamma and Il-2 signaling, consistent with the differences in T-cell abundance (PC2, Fig. 1g). These data highlight the importance of the host immune system in sculpting the transcriptome and provide cautionary context for the analysis of tumor gene expression in immune-compromised hosts. Although there were large differences between clonal gene expression patterns across hosts the clones showed consistent differences in gene expression across all settings, reflective of intrinsic clonal properties, with the biggest variation in gene expression across the clones being related to their position along the epithelial-mesenchymal transition (EMT) axis (PC3, Fig. 1g). In particular, Clone 679 is the most distinct and the most mesenchymal of the clones.

To further characterize the major clones in our tumors, we performed gene set expression analysis using AUCell (Aibar et al., 2017) to identify pathways that are enriched in cells of a specific clonal lineage. Analysis was performed across four independent experiments each with three vehicle-treated animals and for the majority of clones we were able to identify distinct gene expression signatures that were reproducible across animals and experiments (Fig. 1h, Supplementary Table 4, Supplementary Table 5).

### Simultaneous detection of changes in clonal abundance, gene expression, and tumor microenvironment in response to BET bromodomain inhibition with WILD-seq

Having established that we can repeatedly observe the same clonal lineages and their gene expression programs across animals and experiments, we next sought to perturb the system. We chose the BET bromodomain inhibitor JQ1 for our proof-of-principle experiments to assess the ability of the WILD-seq system to simultaneously measure changes in clonal abundance, gene expression and the tumor microenvironment that occur following therapeutic intervention. JQ1 competitively binds to acetylated lysines, displacing BRD4 and thereby repressing transcription at specific loci. A large number of studies have indicated that BET inhibitors may be beneficial in the treatment of hematological malignancies and solid tumors including breast cancer, possibly by inhibiting certain key proto-oncogenes such as MYC (G. Jiang et al., 2020).

Treatment of our 4T1 WILD-seq tumor-bearing mice with JQ1 caused an initial suppression of tumor growth but with only a small overall effect on time to humane endpoint (Fig. 2a). Tumors treated with JQ1 or vehicle alone were harvested at endpoint, dissociated and subjected to single cell sequencing (Fig. 2b). Two independent experiments were performed, each with 3 mice per condition.

**Figure 2.**
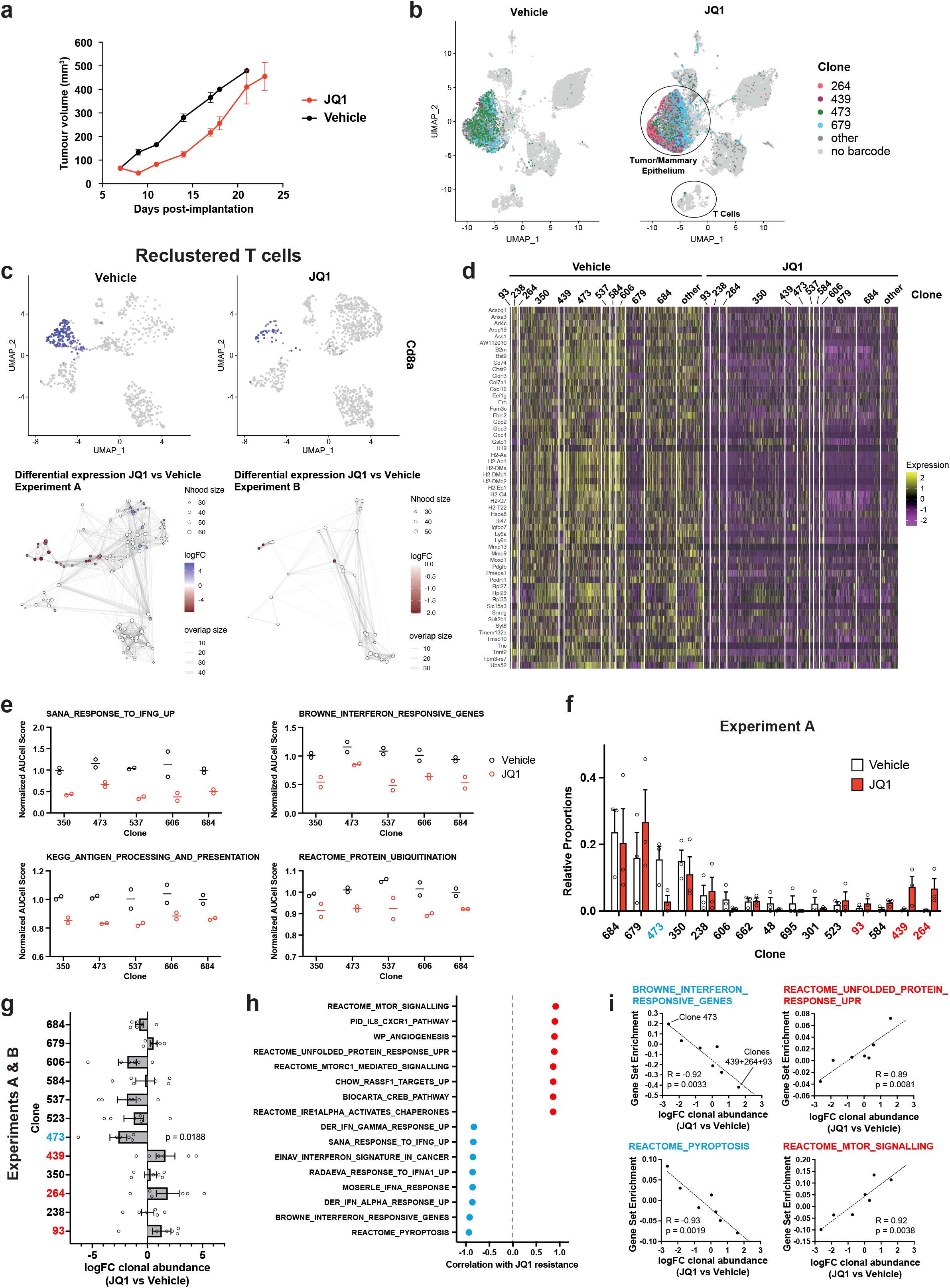
Simultaneous detection of changes in clonal abundance, gene expression, and tumor microenvironment in response to BET bromodomain inhibition with WILD-seq. **a. Tumor growth curves with JQ1 treatment.** 4T1 WILD-seq tumors were treated with the BET bromodomain inhibitor JQ1 or vehicle from 7 days post-implantation until endpoint (n = 4 mice per condition). Data represents mean ± SEM. **b. scRNA-seq of JQ1-treated 4T1 WILD-seq tumors.** UMAP plots of vehicle- or JQ1-treated 4T1 WILD-seq tumors. Combined cells from 2 independent experiments, each with 3 mice per condition are shown. Cells for which a WILD-seq clonal barcode is identified are shown as dark grey or colored spots. Cells which belong to four selected clonal lineages are highlighted. **c. JQ1-treatment results in a reduction in Cd8+ tumor-associated T-cells.** Cells belonging to the T-cell compartment were computationally extracted from the single cell data and reclustered. Upper panels show combined UMAP plots from experiments A and B with Cd8a expression per cell illustrated enabling identification of the Cd8+ T cell cluster. Lower panels show neighborhood graphs of the results from differential abundance testing using Milo (Dann et al., 2021). Colored nodes represent neighbourhoods with significantly different cell numbers between conditions (FDR < 0.05) and the layout of nodes is determined by the position of the neighborhood index cell in the UMAP panel above. Experiments A and B were analyzed separately due to differences in cell numbers. **d. Differential gene expression between JQ1- and vehicle-treated tumor cells.** Single cell heatmap of expression for genes which are significantly and consistently down-regulated across clonal lineages (combined fisher p-value < 0.05 and mean logFC < −0.2 for both experiments). 1600 cells are represented (400 per experiment/condition), grouped according to their clonal lineage. **e. Differential gene set expression between JQ1 and vehicle-treated tumor cells.** Median AUCell score per experiment/condition for selected gene sets. The 5 clonal lineages with the highest representation across experiments are shown. **f. Clonal representation.** Proportion of tumor cells assigned to each clonal lineage in experiment A based on the WILD-seq barcode (n = 3 tumors per condition). Clones which make up at least 2% of the assigned tumor cells under at least one condition are plotted. The most sensitive clone 473 is highlighted in blue and the most resistant clones 93, 439, 264 are highlighted in red. Data represents mean ± SEM. **g. Clonal response to JQ1-treatment.** Log2 fold change in clonal proportions upon JQ1 treatment across experiments A and B. Fold change was calculated by comparing each JQ1-treated sample with the mean of the 3 corresponding vehicle-treated samples from the same experiment. p-value calculated by one-sample t-test vs a theoretical mean of 0. Data represents mean ± SEM. **h. and i. Correlation of JQ1-response with baseline clonal transcriptomic signatures.** Clonal gene set enrichment scores for vehicle-treated tumors were calculated by comparing cells of a specific clonal lineage of interest to all assigned tumor cells within the same experiment. Correlation between these scores and JQ1-treatment response (mean log_2_ fold change clonal proportion JQ1 vs vehicle) was then calculated for each gene set. Selected gene sets with the highest positive or negative correlation values (Pearson correlation test) are shown. A positive correlation indicates a higher expression in resistant clones, whereas a negative correlation indicates a higher expression in sensitive clones. Resistant clonal lineages identified by barcodes 93, 264 and 439 were combined for the purpose of this analysis to have enough cells for analysis within the vehicle-treated samples.

We first explored whether JQ1 had any effect on the tumor microenvironment. The most striking difference we observed was a change in abundance among the cells belonging to the T-cell compartment. To analyze this further, we computationally extracted these cells from the single cell data, reclustered them and performed differential abundance testing using Milo (Fig. 2c). Milo detects sets of cells that are differentially abundant between conditions by modeling counts of cells in neighborhoods of a KNN graph (Dann et al., 2021). When applied to our reclustered T-cells, Milo identified a significant decrease in abundance in cytotoxic T-cells, as identified by their expression of Cd8a and Cd8b1, following JQ1 treatment. A significant change was observed in both of our experiments although the magnitude of the effect was greater in experiment A.

We next examined the effect of JQ1 treatment on the transcriptome of the tumor cells. Differential expression analysis was performed for each clonal lineage and experiment independently. As expected, given its mode of action, we identified significant down-regulation of a wide range of genes with consistent changes across clonal lineages (Fig 2d, supplementary table 6). Among the repressed genes, were a number of genes related to interferon (IFN) signaling and antigen processing and presentation (Fig. 2d, 2e), including GBP2 which is strongly induced by IFN gamma, the MHC class II protein, Cd74, and B2m, a component of the MHC class I complex. JQ1 has previously been reported to directly inhibit transcription of IFN-response genes (Gibbons et al., 2019; Gusyatiner et al., 2021) suggesting this may be due to the direct action of JQ1 within our tumor cells, however JQ1-dependent changes to the tumor microenvironment may also influence these expression pathways.

Our barcoded 4T1 clones showed varied sensitivity to JQ1, with treatment causing reproducible changes to clonal proportions within the tumor (Fig. 2f, 2g, Supplementary Table 2). In particular, one of the most abundant clones, clone 473, is highly sensitive to JQ1 treatment. In contrast, 3 clones were identified as being the most resistant to JQ1 treatment, clones 93, 439 and 264. These clones which together make up less than 5% of the tumor in vehicle treated mice constitute on average 12.8% of the JQ1-treated tumors. To examine baseline transcriptomic signatures of JQ1-sensitivity and resistance, we identified gene sets whose expression in vehicle-treated tumors was highly correlated with response (Figs. 2h, 2i, Supplementary Table 7). Interestingly, interferon signaling which is significantly attenuated in our JQ1-treated tumors is highly correlated with sensitivity to JQ1, suggesting a possible higher dependence of the sensitive clones on these pathways. Conversely resistance is associated with higher levels of unfolded protein response and mTOR signaling consistent with a known role of mTOR-mediated autophagy in resistance to JQ1 (Luan et al., 2019), and cytotoxic synergy between PI3K/mTOR inhibitors and BET inhibitors (Lee et al., 2015; Stratikopoulos et al., 2015).

### Clonal transcriptomic correlates of response and resistance to taxane chemotherapy in the 4T1 mammary carcinoma model

Our studies with JQ1 exemplify the ability of the WILD-seq system to simultaneously measure *in vivo* the effect of therapeutic intervention on clonal dynamics, gene expression and the tumor microenvironment. However, we were interested in using our system to investigate a chemotherapeutic agent currently in use in the clinic. We therefore treated our 4T1 WILD-seq tumor-bearing mice with docetaxel as a representative taxane, a class of drugs which are routinely used to treat triple negative breast cancer patients. As with JQ1, docetaxel treatment resulted in an initial, modest reduction in tumor growth followed by relapse (Fig. 3a). Comparison of vehicle and docetaxel (DTX) treated tumors revealed differential response of clonal lineages to treatment (Figs. 3b, 3c, 3d, Supplementary Table 2) with clone 679 being the most resistant and clone 238 the most sensitive to chemotherapy.

**Figure 3.**
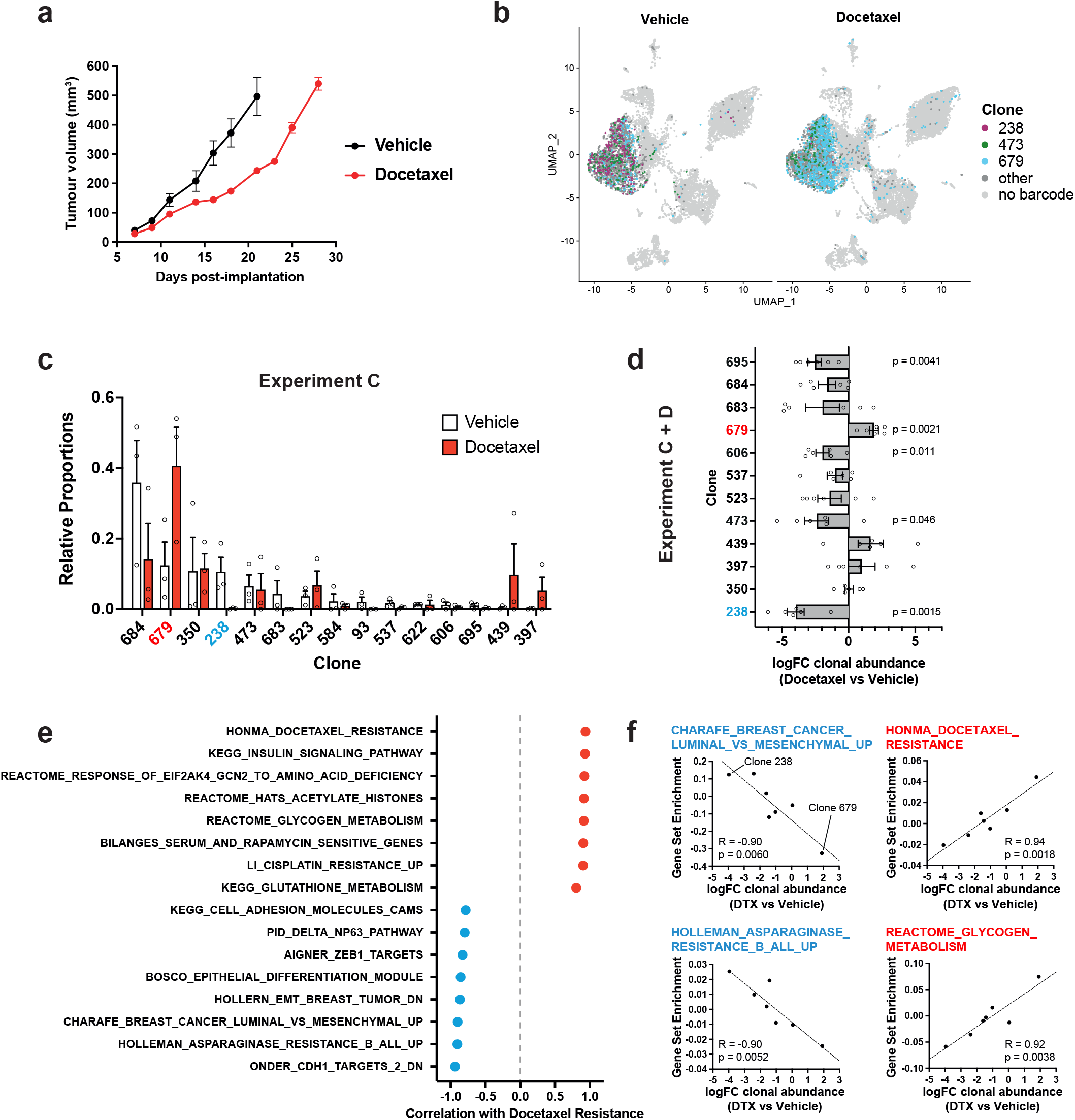
Clonal transcriptomic correlates of response and resistance to taxane chemotherapy in the 4T1 mammary carcinoma model. **a. Tumor growth curves with docetaxel treatment.** 4T1 WILD-seq tumors were treated with docetaxel or vehicle (12.5% ethanol, 12.5% Kolliphor) from 7 days post-implantation for 2 weeks (n = 5 mice per condition). Dosing regimen was 12.5 mg/Kg docetaxel three times per week. Data represents mean ± SEM. **b. scRNA-seq of docetaxel-treated 4T1 WILD-seq tumors.** UMAP plots of vehicle- or docetaxel-treated 4T1 WILD-seq tumors. Combined cells from 2 independent experiments, each with 3 mice per condition are shown. Cells for which a WILD-seq clonal barcode is identified are shown as dark grey or colored spots. Cells which belong to three selected clonal lineages are highlighted. **c. Clonal representation.** Proportion of tumor cells assigned to each clonal lineage in experiment C based on the WILD-seq barcode (n = 3 tumors per condition). Clones which make up at least 2% of the assigned tumor cells under at least one condition are plotted. The most sensitive clone 238 is highlighted in blue and the most resistant clone 679 is highlighted in red. Data represents mean ± SEM. **d. Clonal response to docetaxel-treatment.** Log_2_ fold change in clonal proportions upon docetaxel treatment across experiments C and D. Fold change was calculated by comparing each docetaxel-treated sample with the mean of the 3 corresponding vehicle-treated samples from the same experiment. p-values calculated by one-sample t-test vs a theoretical mean of 0. Data represents mean ± SEM. **e. and f. Correlation of docetaxel-response with baseline clonal transcriptomic signatures.** Clonal gene set enrichment scores for vehicle-treated tumors were calculated by comparing cells of a specific clonal lineage of interest to all assigned tumor cells within the same experiment. Correlation between these scores and docetaxel-treatment response (mean log_2_ fold change clonal proportion docetaxel vs vehicle) was then calculated for each gene set. Selected gene sets with the highest positive or negative correlation values (Pearson correlation test) are shown. A positive correlation indicates a higher expression in resistant clones, whereas a negative correlation indicates a higher expression in sensitive clones.

Correlating the clones’ baseline transcriptomic profiles with response to docetaxel, revealed a major role for EMT in modulating sensitivity and resistance to taxane-based therapy. The 4T1 clones which are most sensitive to docetaxel are characterized by high expression of E-Cadherin regulated genes and low Zeb1 activity consistent with a more epithelial phenotype (Figs. 3e, 3f, Supplementary Table 8). These observations are in agreement with previous studies that have implicated EMT, and its associated endowment of cancer stem cell-like characteristics, as a mechanism of resistance to cytotoxic chemotherapies like docetaxel in cell culture and patients (Bhola et al., 2013; Creighton et al., 2009; Gupta et al., 2009). Resistance to docetaxel was correlated with up-regulation of multiple gene sets (Figs. 3e, 3f, Supplementary Table 8). This included genes whose expression is elevated in non-responders to docetaxel in human breast cancer patients (Honma et al., 2008) demonstrating the relevance of findings arising from this approach. Interestingly, we also identify metabolic reprogramming as a potential mechanism of docetaxel resistance with higher expression of genes related to glycogen and glutathione metabolism being correlated with resistance to docetaxel (Fig, 3e).

### Clonal transcriptomic signatures of response and resistance to taxane chemotherapy in the D2A1 mammary carcinoma model

To explore the general applicability of WILD-seq to other models, we utilized a second triple negative mammary carcinoma model, D2A1-m2 (hereafter referred to as D2A1). Similar to the 4T1 cell line, this line was derived from a mouse mammary tumor in a BALB/c mouse and can be orthotopically implanted into the mammary fat pad of immunocompetent, syngeneic hosts (Jungwirth et al., 2017).

We established a WILD-seq D2A1 clonal pool by transducing the D2A1 cell line with our WILD-seq barcode library. These barcoded cells were orthotopically implanted into a cohort of mice, half of which were treated with docetaxel, while the remaining animals received vehicle alone. Docetaxel treatment caused an initial reduction in tumor growth with subsequent relapse (Fig. 4a). We performed single cell RNA sequencing of three tumors per condition and assigned the tumor cells to a distinct clonal lineage based on the presence of the WILD-seq barcode (Fig. 4b). In total 103 different WILD-seq barcodes were observed *in vivo* with a dramatic shift in relative clonal abundance on docetaxel treatment (Fig. 4d, Supplementary Table 3). Unlike our 4T1 breast cancer model, variation between clonal lineages was no longer dominated by the EMT status of the clones and all clones exhibited a more mesenchymal-like phenotype consistent with the fact that this was a subline of D2A1 selected for its metastatic properties (Fig. 4c). This provides us with a distinct yet complementary system to investigate chemotherapy resistance with the potential to reveal alternative mechanisms than EMT status.

**Figure 4.**
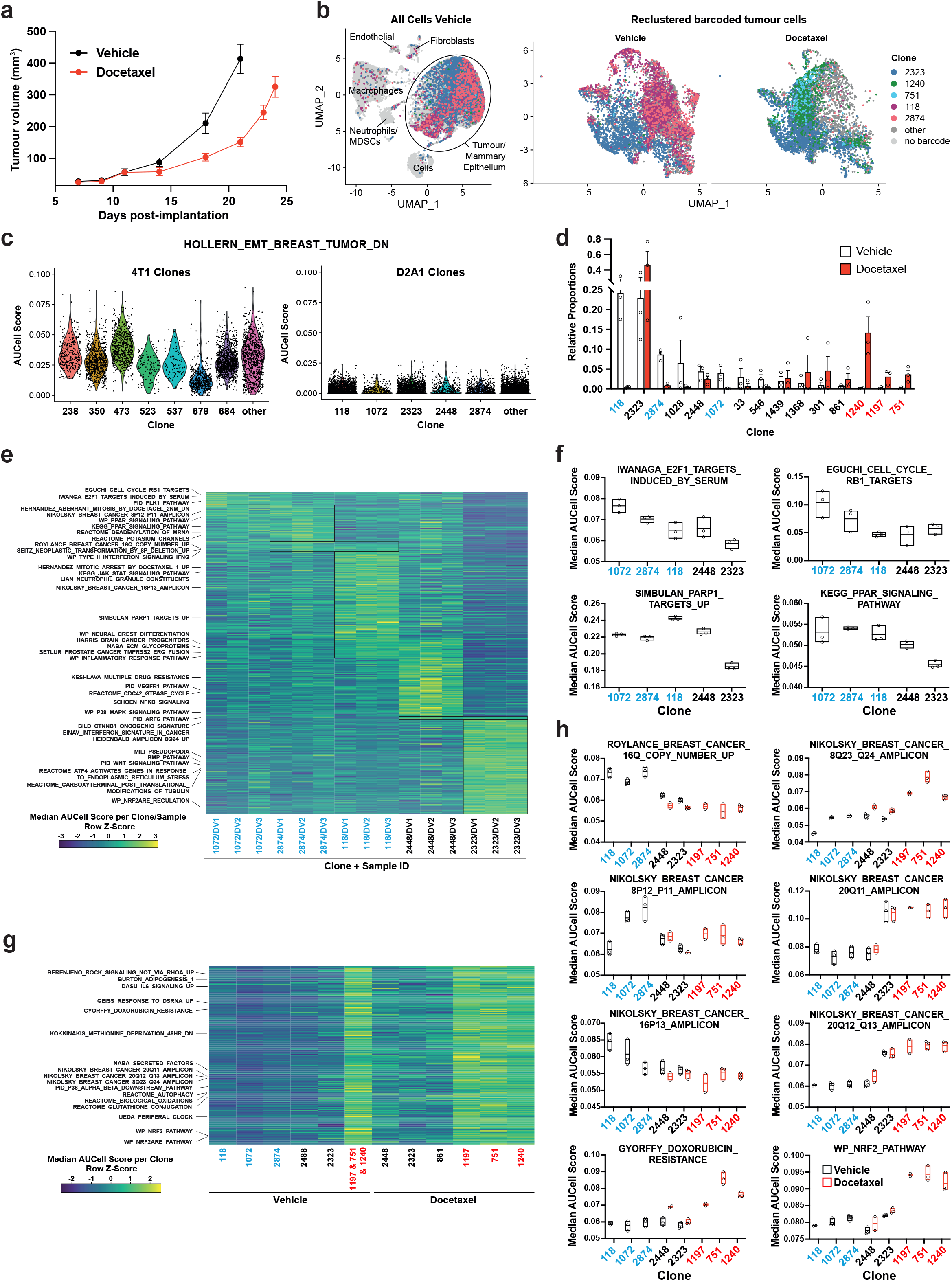
Clonal transcriptomic signatures of response and resistance to taxane chemotherapy in the D2A1 mammary carcinoma model. **a. D2A1 WILD-seq tumor growth curves with docetaxel treatment.** D2A1 WILD-seq tumors were treated with docetaxel or vehicle from 7 days post-implantation for 2 weeks (n = 5 vehicle-treated mice, n = 4 docetaxel-treated mice). Data represents mean ± SEM. **b. scRNA-seq of docetaxel-treated D2A1 WILD-seq tumors.** UMAP plots of vehicle-treated D2A1 WILD-seq D2A1 tumors and reclustered barcoded-tumor cells from vehicle- and docetaxel-treated tumors. Combined cells from 3 mice per condition are shown. Cells for which a WILD-seq clonal barcode is identified are shown as dark grey or colored spots. Cells which belong to five selected clonal lineages are highlighted. **c. Comparison of EMT status of major 4T1 and D2A1 WILD-seq clones.** Violin plot of AUCell scores from vehicle-treated tumor cells generated using the HOLLERN_EMT_BREAST_TUMOR_DN (Hollern et al., 2018) gene set, a set of genes that have low expression in murine mammary tumors of mesenchymal histology. 4T1 WILD-seq clones exhibit varying levels of expression of this geneset whereas D2A1 WILD-seq clones have consistently low levels of expression of these genes. **d. Clonal representation.** Proportion of tumor cells assigned to each clonal lineage based on the WILD-seq barcode (n = 3 tumors per condition). Clones which make up at least 2% of the assigned tumor cells under at least one condition are plotted. The most sensitive clones to docetaxel treatment 118, 2874 and 1072 are highlighted in blue and the most resistant clones 1240, 1197 and 751 are highlighted in red. Data represents mean ± SEM. **e. Clonal transcriptomic signatures from vehicle-treated tumors.** Heatmap of median AUCell scores per sample for each of the five most abundant clones. All gene sets which showed consistent and statistically significant enrichment (combined fisher p-value < 0.01& mean log_2_ enrichment > 0.1) in at least one of these clones are illustrated. **f. Selected gene sets whose expression is associated with sensitivity to docetaxel.** Median AUCell scores per sample for each of the five most abundant clones is plotted. **g. Transcriptomic signatures associated with resistance to docetaxel.** For vehicle-treated tumors, resistant clonal lineages identified by barcodes 1197, 751 and 1240 were combined to have enough cells for analysis. Gene sets with significantly enriched expression in these resistant clones in vehicle-treated tumors were determined (adjusted p-value < 0.01& log_2_ enrichment > 0.1). A heatmap of median AUCell scores per clone, per condition of these resistance-associated gene sets is plotted. **h. Selected gene sets whose expression is enriched or depleted in resistant clones.** Median AUCell scores per clone, per sample are plotted for samples with at least 20 cells per clone. Due to changes in clonal abundance with treatment some clones can only be assessed under vehicle- or docetaxel-treated conditions.

We identified 3 clones which were acutely sensitive to docetaxel, clones 118, 2874 and 1072. Together these constitute on average 37% of the vehicle-treated tumors but only 1.3% of the docetaxel-treated tumors (Fig. 4d). To understand the properties of these clones, we analyzed the baseline gene expression characteristics of clones in vehicle-treated tumors. The gene expression of cells from a clone of interest was compared to all tumor cells to which a WILD-seq barcode could be assigned from the same sample, and clonal signatures identified that were significantly enriched across animals. Specific gene expression signatures were identifiable for all clones analyzed, some of which were unique to a single clone while others overlapped across the sensitive clones (Fig. 4e, Supplementary Table 9, Supplementary Table 10). For example, clone 1072 shows elevated levels of expression of cell cycle related pathways, such as E2F-target genes (Fig 4f), indicating that aberrant cell cycle control in these cells that could increase their susceptibility to an antimitotic cancer drug (Fig. 4f), interestingly high levels of E2F-targets have recently been shown to be associated with response to chemotherapy in breast cancer patients (Sammut et al., 2021).

Three clones robustly increased their relative abundance within the tumor following docetaxel treatment, clones 1197, 751 and 1240, which despite making up less than 1% of the vehicle-treated tumors together constituted more than 20% of the docetaxel-treated tumors (Fig. 4d). Due to the low abundance of cells in vehicle-treated samples, cells belonging to all 3 clones were pooled to analyse their baseline gene expression profiles (Fig. 4g). Among the gene sets differentially expressed between resistant and sensitive clones, were a number of breast cancer amplicons indicating that there may be specific copy number variations associated with these clones (Figs. 4g, 4h). However single cell DNA sequencing data would be required to confirm the presence of specific genetic traits within our clones. Interestingly, gains in 8q24 (Han et al., 2010), 20q11 (Voutsadakis, 2021) and loss of 16q (Höglander et al., 2018) have previously been reported to be associated with resistance to taxane-based chemotherapy in agreement with our findings. Highly upregulated within all 3 of our resistant clones were genes related to the NRF2 pathway, even in the absence of docetaxel treatment (Figs. 4g, 4h). NRF2 activation has been linked to cancer progression and metastasis and has been suggested to confer resistance to chemotherapy (Homma et al., 2009; T. Jiang et al., 2010; Konstantinopoulos et al., 2011; Romero et al., 2017; Shibata et al., 2008; Singh et al., 2006).

### Delineating the contribution of clonal abundance to gene expression changes upon drug treatment

Prior to the advent of single cell sequencing, the majority of studies relied on bulk RNA-seq or microarray analysis of gene expression to interrogate the effect of chemotherapeutic interventions. While informative, these studies cannot differentiate between changes in bulk gene expression that arise due to clonal selection and changes that are induced within a clonal lineage as the result of drug exposure. Even with single cell sequencing, definitive identification of the same clonal population across treatment conditions is impractical. Our method alleviates these difficulties by enabling the direct comparison of clones of the same lineage under different conditions.

To examine the relative contribution of clonal selection and transcriptional reprogramming to changes in gene expression upon chemotherapy, we compared analysis of gene expression within each clone individually to a combined analysis of all pooled tumor cells (Fig. 5, Supplementary Table 11). Consistent with their mode of action, docetaxel had relatively little effect on the transcriptome of individual clones while JQ1 caused substantial changes to the transcriptome predominantly down-regulating gene expression. Genes were identified under all treatments that were altered within the tumor as a whole but as a result of clonal selection rather than intra-clonal changes in gene expression, with the biggest effects being observed with docetaxel treatment in D2A1 tumors, in agreement with this condition inducing the largest changes in relative clonal abundance. To confirm that changes in gene expression detected in bulk tumor analysis but not the clonal analysis could be attributed to differences in clonal sensitivity to chemotherapy, we analyzed baseline expression of these genes across the major clonal populations (Fig. 5b). As expected, we found that genes up-regulated only in bulk tumor analysis had significantly higher expression in clones resistant to chemotherapy (that increase in abundance with treatment) and genes only down-regulated in bulk tumor analysis had significantly lower expression in these resistant clonal lineages.

**Figure 5.**
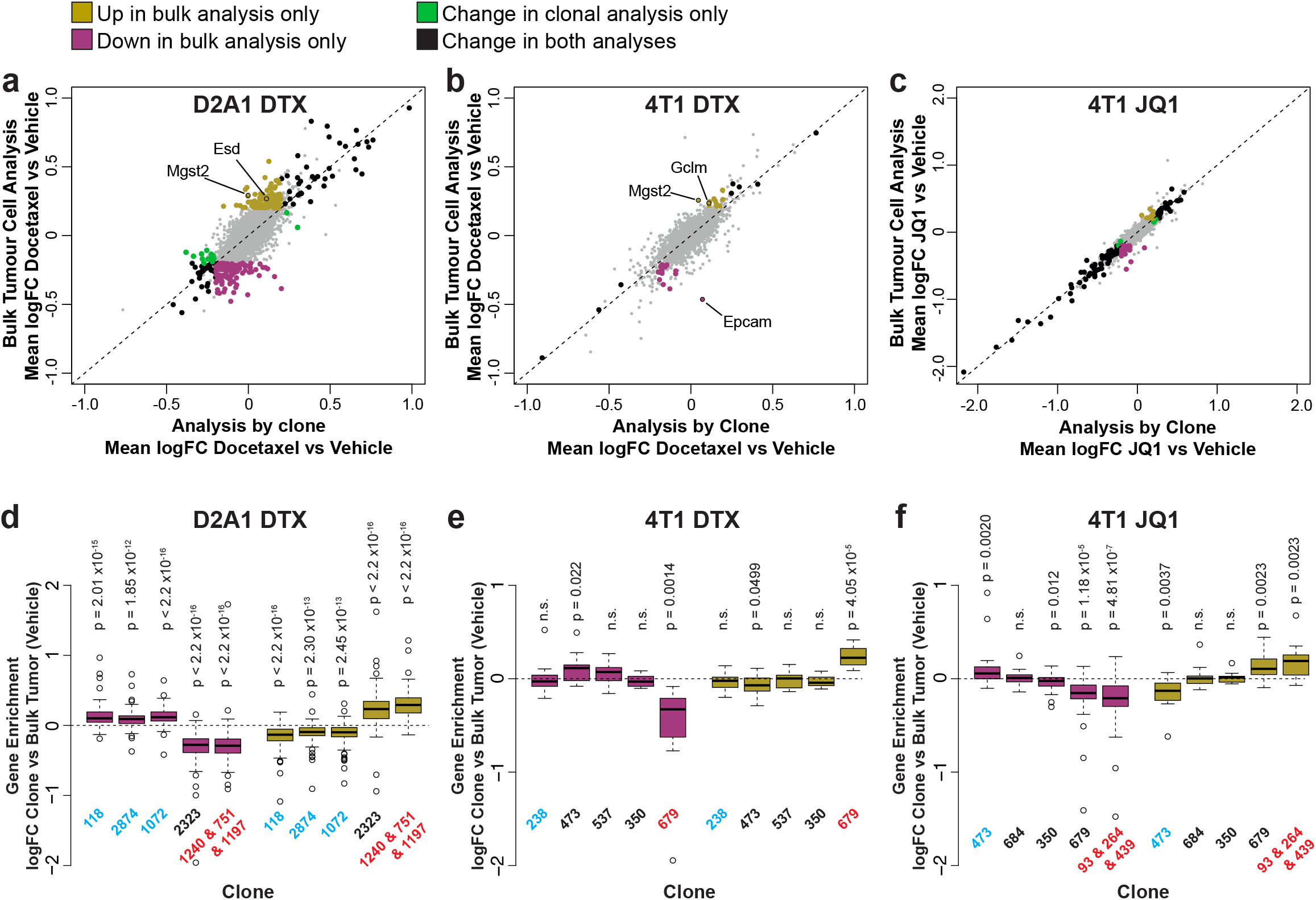
Delineating the contribution of clonal abundance to gene expression changes upon drug treatment. **a. Comparison of differential gene expression analysis in bulk tumor cells and intra-clonal changes in gene expression.** Differential gene expression was performed for all barcoded tumor cells irrespective of clonal lineage comparing chemotherapy-treated and vehicle-treated cells (bulk tumor cell analysis). Alternatively differential gene expression was performed for each individual clone separately and the results combined to identify genes which robustly undergo intra-clonal changes in expression (analysis by clone). Whereas bulk tumor cell analysis will identify changes in overall gene expression due to both changes in clonal abundance and changes within the cells, analysis by clone enables us to delineate exclusively induced cellular changes in gene expression. Log_2_ fold change in expression as determined by each of these analysis methods is plotted. Genes with significant changes in expression with chemotherapy (p-value < 0.05, logFC < −0.2 or > 0.2) are highlighted based on the method under which they were identified. Genes identified as significantly changing by one method only met neither logFC nor p-value cutoffs in the alternative method. **b. Changes in gene expression that are identified by bulk tumor cell analysis only can attributed to changes in clonal abundance.** The expression of genes which were identified as differential expressed after chemotherapy only in the bulk tumor cell analysis was assessed across clonal lineages at baseline. Baseline gene enrichment for each clone was determined as described previously by comparing cells of a specific clonal lineage to all barcoded tumor cells within the same vehicle-treated sample or experiment. Gene enrichment values for all genes with differential expression only in the bulk tumor cell analysis were plotted. As expected, genes down-regulated in bulk analysis have lower expression in resistant clones, whereas genes up-regulated in bulk analysis are enriched in resistant clones. p-values represent a one sample t-test vs a theoretical mean of 0.

Among the genes that change in expression within the tumor as a whole as a result of clonal selection upon docetaxel treatment, we identified a number of genes related to glutathione synthesis and conjugation including Mgst2, Esd and Gclm, that may endow resistant clones with greater ability to resolve reactive oxygen species (ROS) induced by docetaxel (Alexandre et al., 2007). Of note, we also observed that in 4T1 tumors, Epcam was significantly reduced in expression in the bulk tumor but was not changed within the individual clonal populations. This suggests that rather than inducing an EMT within the tumor cells, docetaxel is selecting clones of a pre-existing more mesenchymal phenotype.

### Convergent WILD-seq analysis across models identifies redox defense as a mediator of taxane resistance and amino asparagine deprivation as a means to target resistant clones

To examine if there were any shared mechanisms of taxane resistance across our 4T1 and D2A1 WILD-seq clones, we looked for genes that were enriched in resistant clonal lineages in both models. We identified 47 overlapping resistance genes (Fig. 6a, Supplementary Table 12). These genes were significantly enriched in pathways related to resolution of oxidative stress including the NRF2 pathway and glutathione-mediated detoxification (Fig. 6b).

**Figure 6.**
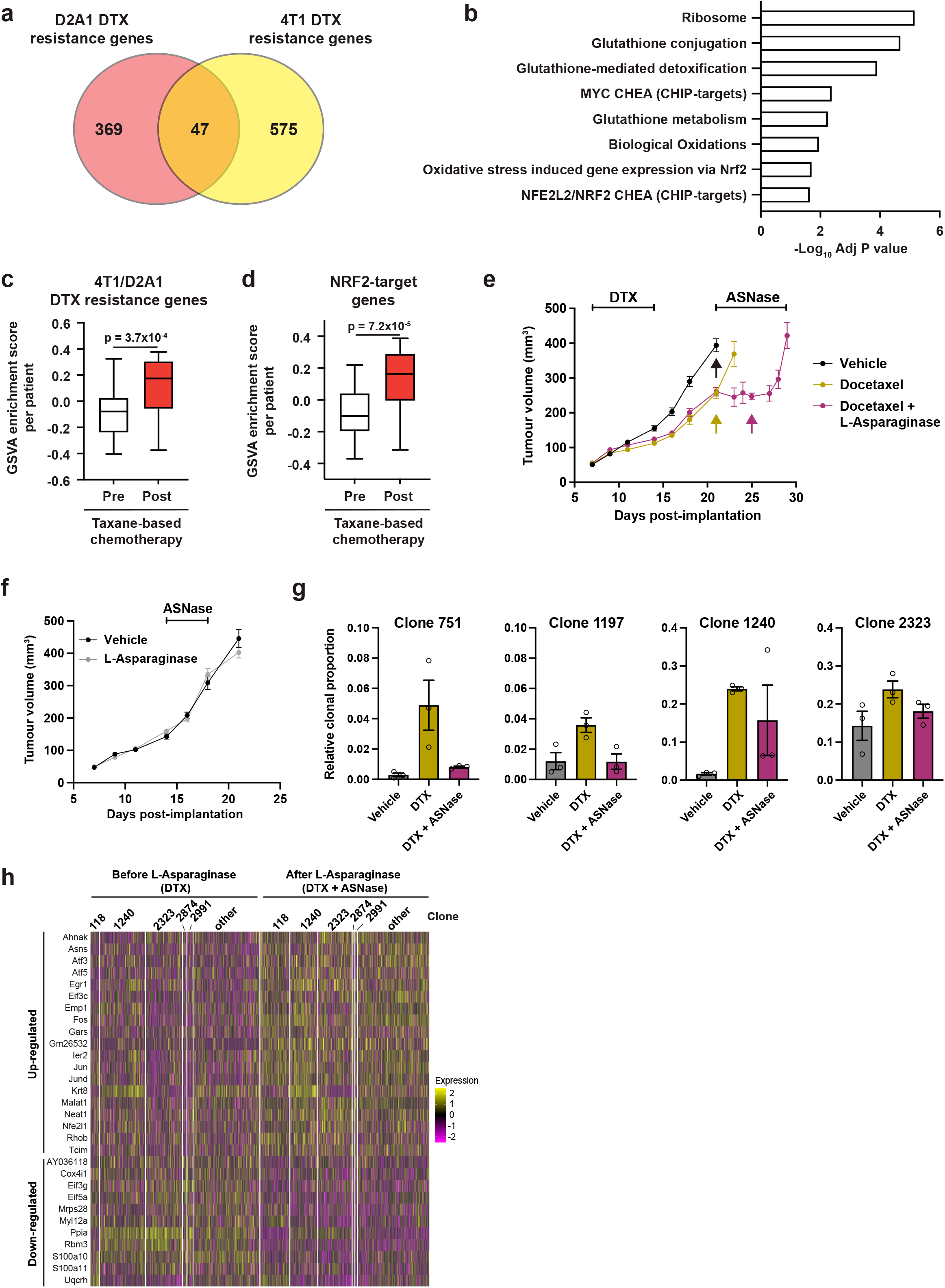
Taxane-resistant clones have elevated NRF2 signaling and are sensitive to asparagine deprivation. **a. Overlap of genes associated with resistance between the D2A1 and 4T1 WILD-seq models.** 4T1 resistance genes were defined as those that were significantly enriched in resistant clone 679 but not in sensitive clone 238 (p < 0.05). D2A1 resistance genes were defined as those that were significantly enriched in combined resistant clones 1240, 751 and 1197 but not in sensitive clones 118, 2874 or 1072 (p < 0.05). In all cases, resistance genes were defined from vehicle treated tumors **b. Gene set enrichment analysis of common resistance genes.** Gene set enrichment was performed using Enrichr for the human orthologs of the 47 common resistance genes identified in Fig. 6a. Adjusted p-values for a subset of significant gene sets are plotted. **c. Expression of our identified resistance genes is increased in human breast tumors following taxane-based chemotherapy.** Expression of our 47 common resistance genes was assessed in human breast cancer samples taken before and after taxane-based neoadjuvant chemotherapy (GSE28844). GSVA enrichment scores for our gene set was calculated for samples from 28 patients for which matched pre- and post-treatment gene expression data were available. Patients received one of three taxane-containing treatment regimens; Regimen A: Epirubicin 90 mg/m^2^-Cyclophosphamide 600 mg/m^2^, 3 cycles bi-weekly and *Paclitaxel* 150 mg/m^2^-Gemcitabine 2500 mg/m^2^, 6 cycles bi-weekly ± weekly Herceptin 4 mg/Kg during the first week, 2 mg/Kg for the remaining 11 cycles. Regimen B: Doxorubicin 60 mg/m^2^-Pemetrexed 500 mg/m^2^, 4 cycles tri-weekly and *Docetaxel* 100 mg/m^2^, 4 cycles tri-weekly. Regimen C: Doxorubicin 60 mg/m^2^-Cyclophosphamide 600 mg/m^2^, 4 cycles tri-weekly and *Docetaxel* 100 mg/m^2^, 4 cycles triweekly. Expression of our common resistance gene set was significantly increased after chemotherapy in human samples. p-value calculated by paired t-test. **d. NRF2-target genes are upregulated in human patients following neoadjuvant chemotherapy.** GSVA enrichment scores for NRF2-target genes (NFE2L2 CHEA consensus CHIP-targets) were calculated for samples from 28 patients in the GSE28844 dataset for which pre- and post-treatment gene expression data were available. p-values calculated by paired t-test. **e. Docetaxel-resistant tumors are collaterally sensitive to L-asparaginase.** D2A1 WILD-seq tumors were treated with 3 doses of 12.5 mg/kg docetaxel (days 7,9,11 post-implantation) and 1 dose of 10 mg/kg docetaxel (day 14 post-implantation). From day 21 mice were treated daily with L-asparaginase. Arrows indicate timepoints of tumor collection for single-cell sequencing. Measurements are combined from 2 independent experiments. Due to sample collection at timepoints indicated the number of animals is reduced beyond this. Vehicle n = 15 mice, docetaxel n = 14 mice (reduced to 5 mice after day 21), docetaxel + L-asparaginase n = 13 mice (reduced to 4 mice from day 25). In addition, 2 mice reached humane endpoint (due to weight loss following docetaxel treatment but prior to administration of L-asparaginase) one in the DTX only arm at day 18 and one in the DTX+L-Asp arm at day 21. Data represents mean ± SEM. **f. L-asparaginase alone does not affect tumor growth.** D2A1 WILD-seq tumors were treated with L-asparaginase or vehicle for 5 consecutive days from day 14 post-implantation. n = 10 mice per condition. Data represents mean ± SEM. **g. Taxane-resistant clones are sensitive to L-asparaginase.** Relative clonal abundance in vehicle-treated (day 21), docetaxel-treated (day 21) and docetaxel and L-asparaginase-treated (day 25) D2A1 WILD-seq tumors is shown for 3 taxane-resistant clones (751, 1197, 1240) and 1 neutral clone (2323). Clonal proportions were calculated from single cell sequencing data of 3 tumors per condition. Data represents mean ± SEM. **h. Gene expression changes in tumor cells after L-asparaginase treatment.** Heatmap for genes which are most significantly and consistently differentially expressed across clonal lineages after treatment with L-asparaginase. 2400 cells are represented (400 per sample), grouped according to clonal lineage.

Importantly, these genes were also enriched in human patients following combined anthracycline and taxane-based therapy, highlighting the potential clinical significance of our findings (Fig. 6c). Gene expression data from a previously published study with paired pre-neo adjuvant chemotherapy (NAC) core needle biopsies and post-chemotherapy surgical samples (Vera-Ramirez et al., 2013) was re-analyzed using GSVA (Hänzelmann et al., 2013) to determine the effect of chemotherapy on a gene set composed of our 47 shared resistance genes (Fig. 6c) as well as NRF2-targets as determined by ChIP enrichment analysis (CHEA) (Lachmann et al., 2010) (Fig. 6d). Expression of both these gene sets was significantly increased after chemotherapy, which our data would suggest is the result of outgrowth of resistant clonal lineages with increased propensity to withstand taxane-induced oxidative stress.

Given these findings, we hypothesized that combining taxane-based chemotherapy with a drug specifically targeting resistant clones with high Nrf2 signaling would provide a highly effective treatment regime. To test this hypothesis, we leveraged the finding that tumors with constitutively active Nrf2, due to mutation in the negative regulator Keap1, have metabolic vulnerabilities that arise from their high antioxidant production (Romero et al., 2017), including dependency of glutamine (Romero et al., 2017) and a general dependency on exogenous non-essential amino acids (NEAA) including asparagine (LeBoeuf et al., 2020). This metabolic dependency can be targeted therapeutically by L-asparaginase (ASNase from *E.coli*), which is used in the clinical management of acute lymphoblastic leukemia (*ALL*)(Batool et al., 2016), and catalyzes the conversion asparagine to aspartic acid and ammonia (Chan et al., 2019).

To ascertain whether docetaxel resistant clones were collaterally sensitive to ASNase, we treated D2A1 WILD-seq tumors initially with docetaxel to select for resistant clones and then began daily treatment with L-asparaginase one week later. This dosing regime was chosen as we found that with the dose of docetaxel used in this study co-administration of the 2 drugs or treatment with ASNase immediately following docetaxel was poorly tolerated. As shown in Figure 6e, treatment with ASNase arrested tumor growth and led to a ~40% increase in time to endpoint (relative to vehicle) in this highly aggressive model, although the tumors did acquire resistance and regrew after approximately one week of treatment. Importantly, ASNase alone had no significant effect on tumor growth, indicative of a docetaxel-induced effect (Fig. 6f). To determine the response of individual clonal lineages to ASNase treatment, we performed single cell sequencing on vehicle treated tumors (day 21), as well as docetaxel treated tumors before the start of ASNase treatment (day 21) and after 4 doses of ASNase (day 25). As before, our docetaxel-resistant clones, 751, 1197 and 1240, which have high levels of Nrf2 signaling all exhibited a dramatic increase in their abundance with docetaxel treatment (Fig. 6g). Excitingly, clones 751 and 1197 were sensitive to ASNase returning to baseline levels. Clone 1240 decreased in abundance in 2 of the 3 mice analyzed so is likely to also be sensitive to ASNase but more data is required to confirm its response. As predicted, our Nrf2-high resistant clones were selectively targeted by amino acid deprivation as other clones such as 2323 were unchanged in their relative abundance (Fig. 6g).

To confirm the mechanism of action of L-asparaginase and identify potential mechanisms of resistance to this drug that might cause the relapse observed, we analyzed the transcriptomic effects of ASNase administration. Genes which consistently changed in expression after ASNase treatment across clonal lineages are shown in Figure 6h. Many of the genes found to be differentially expressed in our tumor cells following L-asparaginase treatment are either directly related to protein synthesis (Eif3c, Gars, Eif3g, Eif5a) or are consistent with changes in gene expression reported in cell lines following amino acid deprivation including Atf5, Atf3, Jun, Fos, Egr1 and Asns (Fu et al., 2011; Pan et al., 2003; Pohjanpelto & Hölttä, 1990; Shan et al., 2010). Of specific interest is the up-regulation of asparagine synthetase (Asns) which catalyzes the *de novo* biosynthesis of L-asparagine from L-aspartate. In acute lymphoblastic leukemia (*ALL*), low levels of ASNS resulting in a dependence on extra-cellular asparagine are considered an important biomarker for L-asparaginase treatment. Moreover, the importance of ASNS overexpression in conferring asparaginase resistance has been well documented and is frequently seen in *ALL* patients that develop drug-resistant forms of the disease following treatment with ASNase (reviewed in (Richards & Kilberg, 2006)). In our experiments, this adaptation to asparaginase is observed across all clones analyzed suggesting a general resistance mechanism and supporting the clinical utility of an Asns inhibitor, if one were to be developed, as third line treatment in this context.

In summary, these data support the notion that WILD-seq can identify causal mechanisms of drug resistance *in vivo*, that can be leveraged to inform new combination therapies. Since the redox defense signatures we identified are detectable in patients after neo-adjuvant chemotherapy (NAC), one can envisage an approach whereby patients receiving NAC have the surgical tumor specimen profiled for NRF2 gene signatures and those with high levels receive a post-operative course of L-asparaginase.

## Discussion

Tumor heterogeneity is thought to underlie drug resistance through the selection of clonal lineages that can preferentially survive therapy. However, identifying the features of such lineages, so that they can be targeted therapeutically, has been challenging due the lack of understanding of their molecular characteristics and the lack of animal models to prospectively test therapeutic interventions and combinations thereof. To overcome these challenges, we utilized WILD-seq, a system that leverages expressed barcodes, population bottle necking, syngeneic mouse models and single cell RNA-seq to link clonal lineage to the transcriptome. Among the existing methods for coupling lineage tracing with single cell transcriptomic profiling, the majority use either lentiviral delivery of a genetic barcode similar to that used here or CRISPR/Cas9-mediate mutations for clonal lineage identification (Biddy et al., 2018; Gutierrez et al., 2021; Quinn et al., 2021; Simeonov et al., 2021; Weinreb et al., 2020). We chose to avoid CRISPR/Cas9-based lineage labeling as induction of DNA damage could have an impact on the transcriptome and the sensitivity of the cells to therapeutic agents (Haapaniemi et al., 2018; L. Jiang et al., 2021). Our approach is unique in that we purposefully bottleneck our clonal population to achieve a balance between maximizing clonal diversity and minimizing variation in clonal representation across replicate animals and experiments. It is this feature that allows us to robustly call clonal gene expression signatures and differential clonal abundance before and after therapeutic intervention and it is this in turn that allows us to identify relevant drug resistance mechanisms *in vivo*.

We find that the abundance of clones in cell culture and *in vivo* differ greatly, with the most abundant clones *in vitro* being lowly represented *in vivo* and vice versa thus providing a cautionary note when analyzing drug response *in vitro*. Moreover, WILD-seq of 4T1 tumors revealed that the relative immune competence of the host profoundly sculpts the transcriptome of clonal lineages and, as exemplified by JQ1, therapeutic interventions can impact the tumor microenvironment and its interaction with tumor cells, effects that would be missed *in vitro* and in immunocompromised hosts. We utilized WILD-seq to analyze sensitivity and resistance to taxane chemotherapy in two syngeneic, triple negative, mammary carcinoma models highlighting both known and new pathways of resistance (Marine et al., 2020). Resistance to cancer therapies can arise due to clonal selection or through adaptive reprograming of the epigenome and transcriptome of individual clones. Our data with docetaxel treatment in 4T1 and D2A1 indicate that over the time frames we have examined clonal selection is the dominant force driving resistance to chemotherapy with gene expression signatures, such as EMT and Nrf2 signaling, being present in clones at baseline that are then selected for during therapy. However, depending on the mode of action of specific drugs, transcriptional reprogramming may also induce therapeutic resistance and such mechanisms can also be effectively identified with the WILD-seq platform. Indeed, up-regulation of Asns, detected across clonal lineages after L-asparaginase provides an example of *de novo* acquisition of a resistance phenotype.

Applying WILD-seq to examine docetaxel response across two TNBC models afforded the opportunity to overlap resistance genes for the same drug across models and remove model-specific effects. These analyses uncovered a critical role for redox defense in docetaxel resistance that also appears to be operative in human breast cancer patients after chemotherapy. Having identified a primary cause of resistance, we next sought to explore the possibility of collateral sensitivity. Collateral sensitivity, first described for antibiotics (Imamovic & Sommer, 2013; Pluchino et al., 2012; Roemhild & Andersson, 2021) is the phenomenon by which resistance to one drug comes at the cost of sensitivity to a second drug. In the context of cancer and taxanes, collateral sensitivity has the distinct advantage over other therapeutic strategies of maintaining the initial first line therapy and only modifying subsequent therapies. We took advantage of previous findings linking constitutive Nrf2 signaling, via Keap1 loss, to a dependency on exogenous non-essential amino acids (LeBoeuf et al., 2020) and thereby sensitivity to L-asparaginase. Application of L-asparaginase led to an initial cessation of tumor growth followed by regrowth 6 days later. WILD-seq of docetaxel treated tumors before and after L-asparaginase treatment confirmed the specific suppression of Nrf2 high clones and also revealed a compensatory, clone agnostic, up-regulation of asparagine synthetase (Asns), which likely drives relapse in these tumors given the importance of ASNS to L-asparaginase resistance in *ALL* (Richards & Kilberg, 2006). Interestingly, we have previously shown that asparagine bioavailability regulates EMT and metastatic progression in breast cancer models (Knott et al., 2018). Thus, asparagine deprivation, which has not been extensively explored in breast cancer, may present multiple benefits to patients and the utility of L-asparaginase, a clinical stage drug, in this setting warrants further investigation.

This study highlights the challenges of tackling tumor heterogeneity therapeutically. Even though we can effectively suppress the induction of docetaxel resistant clones by administration of L-asparaginase the tumors still adapt to this intervention and regrow, most likely due to transcriptionally shifting their metabolism towards *de novo* asparagine synthesis. Nevertheless, hope still remains since there are only three avenues by which cells can supply themselves with asparagine (1) uptake of extra-cellular asparagine which is effectively shut-off by ASNase (2) *de novo* synthesis through Asns or (3) catabolism of existing proteins. If we could effectively force tumors to depend on synthesis through Asns, we could then deprive them of that additional dependency if Asns-directed therapeutics were to be developed. This concept of steering clonal evolution with drugs towards a predictable and irreconcilable, therapeutically targetable, dependency may provide a general approach to achieving durable therapeutic responses for which tractable models of tumor evolution, such as those described here, are essential predictive components.

## Acknowledgments

We would like to thank all members of the Hannon and IMAXT labs at the CRUK Cambridge Institute for valuable discussions and advice, in particular Clare Rebbeck for assistance with home office animal licensing. We would also like to thank CRUK Cambridge Institute’s core facilities, in particular members of the genomics core for assistance with sequencing library preparation and sequencing, members of the Biological Resource Unit (BRU) for animal husbandry and members of the flow cytometry core for assistance with cell sorting. This work was supported by Cancer Research UK [C14303/A21143, C14303/A19926, C14303/A19927, C9545/A24042]. G.J.H. is a Royal Society Wolfson Research Professor.

## Author contributions

Conceptualization- SAW, KS, IGC, GJH. Data Curation- SAW, IGC, KS. Formal Analysis- SAW, IGC, KS. Funding Acquisition- KS, GJH. Investigation- SAW, IGC, KK. Methodology- SAW, IGC, KS. Project Administration- SAW, KS, IGC, GJH, Software- SAW, KS. Supervision- KS, IGC, GJH. Visualization- SAW, KS, IGC. Writing- Original Draft- SAW, KS, IGC. Writing- reviewing and editing- SAW, KS, IGC, GJH.

## Data availability

Single cell RNA-Seq data are being deposited in the gene expression omnibus (GEO) and will be made available upon publication. All other data are available from the corresponding authors upon reasonable request.

## Methods

### Cell lines and culture

The mouse mammary tumor cell lines 4T1 (ATCC) and D2A1-m2 (kind gifted from Clare Isacke’s lab) and the 293FT (Thermo Fisher Scientific) packaging cell line for virus production were cultivated in DMEM high glucose (Gibco), supplemented with 10% heat-inactivated fetal bovine serum (Gibco) and 50 U/mL penicillin-streptomycin (Gibco).

### Virus production

The WILD-seq library was packaged using 293FT lentivirus packaging cells. Cells were plated on 15 cm adherent tissue culture plates (Corning) one day before transfection at a confluency of ~70%. Lentiviral particles were produced by co-transfecting 293FT cells with the transfer plasmid and standard third-generation packaging vectors pMDL (12.5 μg), CMV-Rev (6.25 μg) and VSV-G (9 μg) using the calcium-phosphate transfection method (Invitrogen). The transfection mixture was added to the packaging cells along with 100 mM chloroquine (Sigma-Aldrich). After 16-18 h, media was replaced for fresh growth media. Viral supernatant was collected 48 h after transfection and filtered through a 45μm filter. The viral supernatant was applied directly to cells or stored at 4°C for short-term storage or −80°C for long-term storage. When necessary, virus was concentrated using ultracentrifugation. Lentiviral titre was determined by serial dilutions and measurements of fluorescence via flow cytometry.

### WILD-seq library design and cloning

The pHSW8 lentiviral backbone was constructed using a four-way Gibson Assembly (NEB) by inserting a reverse expression cassette, consisting of a PGK promoter, the zsGreen ORF, a cloning site for high-diversity barcode libraries and a synthetic polyA signal, into an empty pCCL-c-MNDU3-X backbone (#81071 Addgene). To generate the WILD-seq library, a barcode cassette was introduced at the cloning site within the pHSW8 lentiviral backbone, using PCR (Q5 High-Fidelity DNA Polymerase, NEB) and Gibson Assembly (NEB), such that it is expressed within the 3’UTR of the zsGreen transcript.

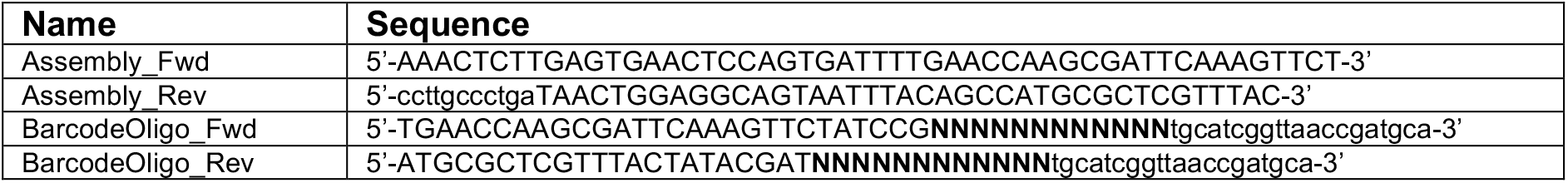

The barcode library was designed by generating 12 nt variable sequences using the R package DNABarcodes (Buschmann, 2017) and a set Hamming distance of 5. The resulting pool of sequences was then purchased as a custom oligo pool (Twist Bioscience). Reverse complement oligos (BarcodeOligo_Fwd/Rev) each containing a specific PCR handle, a 12-bp variable region and 20-bp constant linker were annealed and amplified by PCR for 20 cycles (using Assembly_Fwd/Rev primers). The amplified barcode library was column purified (Gel extraction kit, Qiagen) and the vector backbone was prepared by digestion with SwaI (NEB). WILD-seq barcodes were inserted into the lentiviral vector backbone through Gibson Assembly (NEB), concentrated and transformed into 10b electrocompetent *E.coli* cells (NEB).

### Bottlenecking strategy and characterisation of WILD-seq pools

4T1 or D2A1-m2 cells were infected with WILD-seq library at low MOI (~ 0.2-0.3). Two days after infection, the desired number of zsGreen positive cells, ranging from 10 to 1250 cells, were collected and cultured for two weeks to allow for the pool of clones to stabilize. Different pooling strategies were tested, the ultimate WILD-seq pool was generated from three independent pools each established from 250 sorted cells, maintained separately and mixed in equal proportions immediately prior to injection.

### Library complexity analysis

WILD-seq barcodes of the lentiviral library were amplified using a one-step PCR protocol. 1 ng plasmid was used as template in four separate PCR reactions to account for PCR biases and errors. All reactions were pooled, concentrated and purified on a column and then sequenced on one lane of HiSeq4000. Reads that contained the WILD-seq barcode motif were identified and extracted from the FASTQ files. Detected WILD-seq barcode were filtered based on a 90^th^ percentile cut-off. The resulting whitelist was further filtered for barcodes that contain the common linker region.

### Whitelist generation of WILD-seq barcodes

To generate a comprehensive whitelist of expressed barcodes in each pool, RNA was extracted from WILD-seq transduced cells (High Pure RNA isolation kit, Roche) and reverse transcribed using the Superscript IV reverse transcription kit (Invitrogen) and a target site-specific primer with a unique molecular identifier (UMI) and an Illumina sample index. cDNA was amplified by PCR (Q5 High-Fidelity DNA Polymerase, NEB) using primers (RTWhitelist_Fwd/Rev) containing Illumina-compatible adapters. Alternatively, 1 μg of gDNA was extracted from WILD-seq transduced cells (Blood&Cell Culture DNA Kit, Qiagen) and the barcode amplified by PCR using primers containing Illumina-compatible adapters (gDNAWhitelist_Fwd/Rev). PCR products were purified via gel extraction (Qiagen) and quantified by Qubit. The library was sequenced on an Illumina MiSeq with a custom sequencing primer for Read1 (CustomRead1).

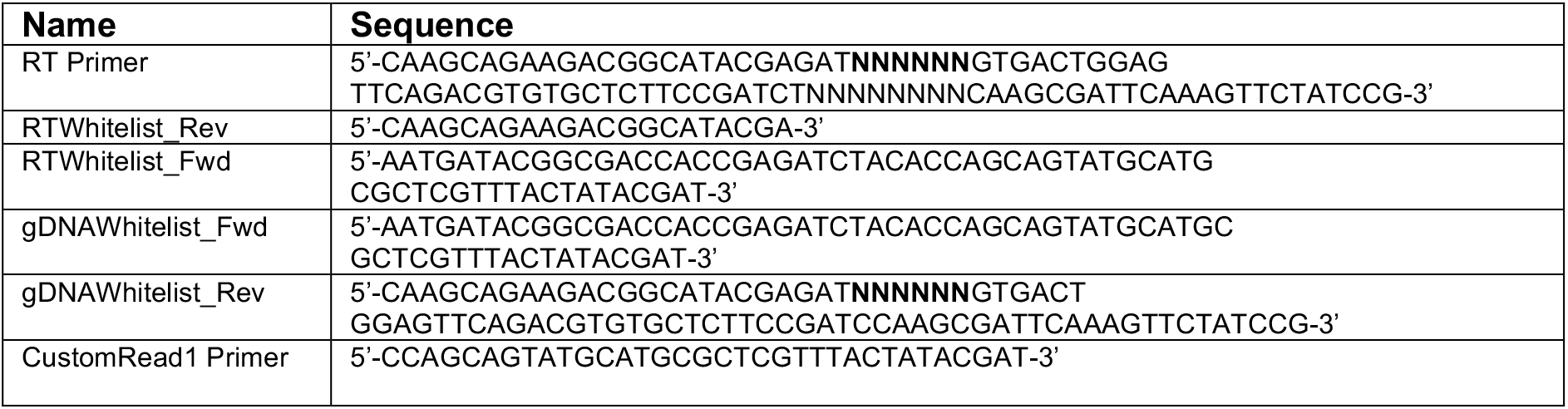

Reads from the RT-PCR barcode library that contained the WILD-seq barcode motif were identified and the number of unique UMIs supporting each barcode was calculated. If barcode sequences amplified from gDNA were also available an additional filtering step was included and any barcodes not also detected in the gDNA library excluded from the whitelist. Based on UMI counts, the top 90^th^ percentile of detected barcodes were taken and collapsed for PCR and sequencing errors using hierarchical clustering and combining sequences with a Hamming distance less than 5.

### Single cell library preparation

Tumor tissues were collected, minced and dissociated using the gentleMACS Octo Dissociator (Miltenyi Biotec) and the relevant kit (Tumor Dissociation Kit mouse). Tissues were process into single cell suspensions following manufacturer’s instructions and filtered through 70 μm filters (Miltenyi) to remove any remaining larger particles from single cell suspension after dissociation. The cell suspension was concentrated and filtered again through a 70 μm filter. Three million live cells were sorted based on live-dead staining with propidium iodide to remove dead cells and debris, pelleted and resuspended in 1 mL phosphate-buffered saline with 0.04% bovine serum albumin (Sigma Aldrich). Cells were counted with a hemocytometer to ensure accurate concentration. The final single cell suspension was diluted as required and NGS libraries were prepared using Chromium Single Cell 3’ Reagent Kit (v3.1 Chemistry Dual Index, user guide reference: CG000315) with no modifications.

### Enrichment library preparation

To enrich for WILD-seq barcodes, the amplified cDNA libraries were further amplified with WILD-seq-specific primers containing Illumina-compatible adapters and sample indices:

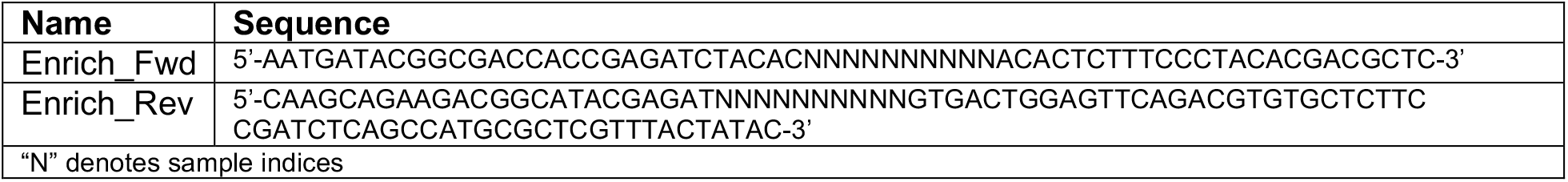

1 μL amplified cDNA library was used as template in a 29-cycle PCR reaction using KAPA HiFi HotStart ReadyMix (Roche). To avoid possible PCR-induced library biases, six reactions were run in parallel. All reactions were combined, purified by columns (Gel purification kit, Qiagen) and quantified by Qubit. Gene expression libraries and barcode enrichment libraries were pooled in an approximately 10:1 molar ratio and libraries were sequenced on the NovaSeq platform (Illumina).

### Animals and *in vivo* dosing

All mouse experiments were performed under the Animals (Scientific Procedures) Act 1986 in accordance with UK Home Office licenses (Project License # PAD85403A) and approved by the Cancer Research UK (CRUK) Cambridge Institute Animal Welfare and Ethical Review Board. Female six to eight week-old BALB/c were purchased from The Charles River Laboratory. 60,000 tumor cells were resuspended in 50 μL of a 1:1 mixture of PBS and growth-factor reduced Matrigel (Corning). All orthotopic injections were performed into the fourth mammary gland. Primary tumor volume was measured using the formula *V=0.5(LxW^2^*), in which *W* is the with and *L* is length of the primary tumor.

Tumor-bearing mice were treated with either vehicle or with different drugs from seven days post transplantation. All drugs were administered via intraperitoneal injection. For JQ1 treatment, animals were dosed 75 mg/kg JQ1 (dissolved in DMSO and diluted 1:10 in 10%β-cyclodextrin) 5 days/week (5 consecutive days followed by 2 days off) until tumors reached endpoint. For docetaxel treatment, animals were dosed at 12.5 mg/kg docetaxel (dissolved in 1:1 mixture of ethanol and Kolliphor and diluted 1:4 in saline) 3 times/week, except when L-asparaginase was to be administered concurrently and then the dose was reduced to 10 mg/kg. For L-asparaginase treatment, mice were administered 100 μL of 60 U L-asparagine (Abcam) diluted in saline. Vehicle-treated mice were sacrificed 21 days post tumor transplantation and treated animals were sacrificed when tumor volumes reached that of vehicle treated animals at 21 days unless otherwise stated.

### scRNA-seq analysis

scRNA-seq libraries generated by the 10X Chromium platform were processed using CellRanger version 3.0.1. Reads were aligned to a custom reference genome that was created by adding the sequence of the zsGreen-WILD-seq barcode transgene as a new chromosome to the mm10 mouse genome. The gene expression matrices generated were then analyzed with the Seurat R package (Stuart et al., 2019) using a standard pipeline. Briefly, datasets were first filtered based on the number of unique genes detected per cell (typical accepted range 200-10000 genes) and the percentage of reads that map to the mitochondrial genome (< 12 %). Reads which mapped to the zsGreen-WILD-seq barcode transgene were removed from the count matrix to prevent these driving cell clustering. Normalisation was performed using sctransform, including cell cycle regression. Differential abundance of cell subtypes was performed using Milo (Dann et al., 2021).

### Clonal barcode assignment to single cell data

#### Extraction of WILD-seq barcodes from scRNA-seq data

Reads mapping to the zsGreen-WILD-seq barcode transgene and containing the full barcode sequence (20nt constant linker with a 12 nt variable region on either side) were extracted from the BAM file produced by Cell Ranger and mapped using Bowtie to a whitelist of barcodes expressed in the WILD-seq cell pool. A WILD-seq clonal barcode was assigned to a cell if there were at least 2 independent reads which matched the barcode to the cell and more than 50% of barcode mapped reads from the cell supported the assignment.

#### Extraction of WILD-seq barcodes from PCR enrichment data

Reads from the PCR barcode enrichment were processed separately using the UMI-tools to extract 10X cell barcodes and UMIs from the raw read files. The sequence corresponding to the full barcode sequence (20nt constant linker with a 12 nt variable region on either side) was extracted from each read and then mapped to the WILD-seq clonal barcode whitelist using Bowtie. A WILD-seq clonal barcode was assigned to a cell if there were at least 10 UMIs which matched the barcode to the cell and at least twice as many UMIs supporting this assignment compared to the next best.

#### WILD-seq barcode assignment

The WILD-seq clonal barcode assignment from these 2 pipelines was then compared. If the assignment from the transcriptomic analysis and the PCR enrichment analysis were in agreement the barcode was assigned. On the rare occasion the assignment didn’t match a clonal barcode was not assigned. If a cell was assigned a WILD-seq barcode by only one method, a further more stringent filtering step was included. For WILD-seq barcodes assigned only from the 10X scRNA-seq dataset but not the PCR-enrichment, the minimum number of UMIs required to support the assignment was increased to 5 and for WILD-seq barcodes assigned only from the PCR-enrichment but not the 10X scRNA-seq dataset, the minimum number of UMIs required to support the assignment was increased to 30.

### Differential gene expression

Differential gene expression was determined using the FindMarkers function in Seurat with a Wilcoxon rank sum test to identify differentially expressed genes. For differential expression of groups of genes, we used the AUCell R package (Aibar et al., 2017) which enables analysis of the relative expression of a gene set (i.e. gene signature or pathway) across all the cells in single-cell RNA-seq data using the “Area Under the Curve” (AUC) to calculate the enrichment of the input geneset within the expressed genes for each cell. An AUCell score was calculated for each tumor cell for every gene set in the MSigDB C2 collection (Liberzon et al., 2011; Subramanian et al., 2005) that contained more than 20 genes with detectable expression in our data. AUCell scores were compared across clones or conditions using a Wilcoxon rank sum test and p-values were adjusted for multiple comparison using the Benjamini-Hochberg correction method.

To generate baseline transcriptomic signatures for each clone in vehicle-treated tumors, comparisons were made between the clone of interest and all assigned tumor cells from the same sample (in the case of D2A1 tumors) or the same experiment (in the case of 4T1 tumors). Samples/experiments were included if they contained at least 20 cells assigned to the clone of interest. To define consistently enriched/depleted signatures, p-values from comparisons within each sample/experiment were combined using the Fisher’s method.

### Patient data analysis

Microarray gene expression data was downloaded from GSE28844 (Vera-Ramirez et al., 2013). A single probe for each gene was selected based on the highest median expression. Gene set expression per patient sample was calculated using GSVA (Hänzelmann et al., 2013).

## Description of supplementary tables

**Supplementary Table 1. Overview of single cell RNA-seq samples generated.**

**Supplementary Table 2. Number and proportion of tumor cells assigned to each clonal barcode for all 4T1 WILD-seq sample.**

**Supplementary Table 3. Number and proportion of tumor cells assigned to each clonal barcode for all D2A1 WILD-seq sample.**

**Supplementary Table 4. 4T1 WILD-seq baseline gene enrichment signatures for major clones.** Differential gene expression analysis was performed for each clone by comparing cells from a clonal lineage of interest to all assigned tumor cells within the same experiment. Only vehicle-treated samples were included in the analysis. Experiments were included in the analysis if they contained at least 20 cells assigned to the clone and clones were analyzed if they were represented by at least 20 cells in at least 3 of the 4 experiments. Differential gene expression was performed using Seurat FindMarkers function and Wilcoxon Rank Sum test. Fisher’s method was used to combine p-values from separate experiments. Analysis for each clone is provided as a separate tab.

**Supplementary Table 5. 4T1 WILD-seq baseline gene set enrichment signatures for major clones.** Differential gene set expression analysis was performed for each clone by comparing cells from a clonal lineage of interest to all assigned tumor cells within the same experiment. All gene sets from the Molecular Signatures Database C2 curated gene set collection were included in the analysis that contained more than 20 genes detectable in our single cell data. Only vehicle-treated samples were included in the analysis. Experiments were included in the analysis if they contained at least 20 cells assigned to the clone and clones were analyzed if they were represented by at least 20 cells in at least 3 of the 4 experiments. Gene set expression analysis was performed using AUCell and differential expression was calculated using Wilcoxon Rank Sum test. Tables show median AUCell score per experiment for each gene set, enrichment in AUCell score relative to all assigned tumor cells within the same experiment (log2(median AUCell score clone of interest/median AUCell score all clones)) and adjusted p-value from Wilcoxon Rank Sum test of AUCell scores from clone of interest vs AUCell scores from all assigned tumors cells from the same experiment. Fisher’s method was used to combine p-values from separate experiments. Analysis for each clone is provided as a separate tab. A final tab ‘Data_for_Fig1h’ provides the matrix of AUCell enrichment values used for the heatmap plotted in figure 1h compiled from individual analyses.

**Supplementary Table 6. Differential expression analysis JQ1 vs Vehicle.** Differential gene expression analysis was performed by comparing cells from the same clonal lineage treated with JQ1 or vehicle within the same experiment. Five clones were included in the analysis (clones 350, 473, 537, 606 and 684) for which there were at least 20 cells per condition across both experiments. Fisher’s method was used to combine p-values from different clones within the same experiment. Gene level differential expression was performed using Seurat FindMarkers function and Wilcoxon Rank Sum test. These data are provided under the ‘FindMarkers_JQ1vsVeh’ tab. Gene set level differential expression was performed using AUCell and differential expression was calculated using Wilcoxon Rank Sum test. These data are provided under the ‘AUCell_JQ1vsVeh’ tab. The ‘Median_norm_AUCell_Scores’ tab provides a summary of the median normalised AUCell scores for each clone, condition and experiment used in the preparation of figure 2e. Normalization to enable comparison across separate experiments was performed by dividing by the median AUCell score for all vehicle-treated tumor cells assigned to any clonal lineage from the same experiment.

**Supplementary Table 7. Correlation of clonal gene expression with JQ1 response.** To determine genes and gene sets whose expression correlates with JQ1 response, the correlation between baseline gene and geneset enrichment values for the major clones as defined in supplementary tables 4 and 5 and the log fold change in clonal abundance between JQ1 and vehicle-treated samples was calculated using the Pearson correlation test. The Pearson correlation coefficient is provided for each gene and gene set.

**Supplementary Table 8. Correlation of clonal gene expression with docetaxel response.** To determine genes and gene sets whose expression correlates with docetaxel response, the correlation between baseline gene and geneset enrichment values for the major clones as defined in supplementary tables 4 and 5 and the log fold change in clonal abundance between JQ1 and vehicle-treated samples was calculated using the Pearson correlation test. The Pearson correlation coefficient is provided for each gene and gene set.

**Supplementary Table 9. D2A1 WILD-seq baseline gene enrichment signatures for major clones.** Differential gene expression analysis was performed for each clone by comparing cells from a clonal lineage of interest to all assigned tumor cells within the same sample. Only vehicle-treated samples were included in the analysis. Clones were included in the analysis if there were at least 20 cells assigned to that clone in all three vehicle samples (DV1, DV2 and DV3). Differential gene expression was performed using Seurat FindMarkers function and Wilcoxon Rank Sum test. Fisher’s method was used to combine p-values from separate samples. Analysis for each clone is provided as a separate tab. In addition, analysis is included for the combined resistant clones 751 1197 and 1240. Due to their low representation in vehicle-treated samples cells assigned to these clones from all three vehicle-treated samples were combined for gene expression analysis and compared to all assigned tumor cells from the three samples.

**Supplementary Table 10. D2A1 WILD-seq baseline gene set enrichment signatures for major clones.** Differential gene set expression analysis was performed for each clone by comparing cells from a clonal lineage of interest to all assigned tumor cells within the same sample. All gene sets from the Molecular Signatures Database C2 curated gene set collection were included in the analysis that contained more than 20 genes detectable in our single cell data. Only vehicle-treated samples were included in the analysis. Clones were included in the analysis if there were at least 20 cells assigned to that clone in all three vehicle samples (DV1, DV2 and DV3). Gene set expression analysis was performed using AUCell and differential expression was calculated using Wilcoxon Rank Sum test. Tables show median AUCell score per sample for each gene set, enrichment in AUCell score relative to all assigned tumor cells within the same experiment (log2(median AUCell score clone of interest/median AUCell score all clones)) and adjusted p-value from Wilcoxon Rank Sum test of AUCell scores from clone of interest vs AUCell scores from all assigned tumors cells from the same sample. Fisher’s method was used to combine p-values from separate samples. Analysis for each clone is provided as a separate tab. In addition, analysis is included for the combined resistant clones 751 1197 and 1240. Due to their low representation in vehicle-treated samples cells assigned to these clones from all three vehicle-treated samples were combined for gene expression analysis and compared to all assigned tumor cells from the three samples. The tab ‘Data for Fig4e and 4f’ provides the matrix of median AUCell scores used for the heatmap plotted in figure 4e compiled from individual analyses. The tab ‘Data for Fig4h’ provides median AUCell scores per sample for clones of interest for all samples and conditions where at least 20 cells per clone were present. Selected data from this table was plotted in figure 4h.

**Supplementary Table 11. Comparison of differential gene expression analysis in bulk tumor cells and intra-clonal changes in gene expression.** For each treatment condition (docetaxel/D2A1, docetaxel/4T1 and JQ1/4T1) differential expression analysis was performed between barcoded tumor cells from drug-treated and vehicle-treated animals from the same experiment. Analysis was performed either by using cells from a single clonal lineage (analysis by clone) or all barcoded tumor cells irrespective of clonal lineage (bulk tumor cell analysis). Differential gene expression was performed using Seurat FindMarkers function and Wilcoxon Rank Sum test. Log2 fold change and adjusted p-value are provided for each comparison. For the analysis by clone, the mean logFC of all individual clonal comparisons is given (mean.logFC.clonal) and Fisher’s method was used to combine p-values (fisher.combined.pvalue.clonal). Genes were classified as significantly changed in clonal analysis only, bulk analysis only or both analysis methods based on significance cutoffs of p-value < 0.05 and logFC < −0.2 or > 0.2. Genes identified as significantly changing by one method only met neither logFC nor p-value cutoffs in the alternative method. For analysis of WILD-seq 4T1 data, analysis was performed separately for the 2 experiments and genes had to meet significance cutoffs in both experiments.

**Supplementary Table 12. Overlap of docetaxel resistance markers in 4T1 and D2A1 cell lines.** 4T1 resistance genes were defined as those that were significantly enriched in resistant clone 679 but not in sensitive clone 238 (p < 0.05). D2A1 resistance genes were defined as those that were significantly enriched in combined resistant clones 1240, 751 and 1197 but not in sensitive clones 118, 2874 or 1072 (p < 0.05). Overlap of these lists revealed 47 common genes. These are listed along with their human orthologs.

**Supplementary Table 13. Number and proportion of tumor cells assigned to each clonal barcode for docetaxel and L-asparaginase combination experiment.**

**Supplementary Table 14. Differential expression analysis for L-Asparaginase treatment.**

Differential gene expression analysis was performed by comparing cells from the same clonal lineage between each DTX+Asp sample and the combined DTX only samples. To ensure there were sufficient cells across all samples, five major clones (118, 1240, 2323, 2874 and 2991) were included in the analysis. Differential expression analysis was performed using Seurat FindMarkers function and Wilcoxon Rank Sum test. Fisher’s method was used to combine p-values from different clones within the same comparison. When selecting genes of interest, mean fold change between DTX+Asp samples and vehicle (also calculated on a per clone basis using abundant clones) was used as an additional cutoff and is included in the table. The most significantly and consistently differentially expressed genes are indicated in the final column ‘Meets.cutoffs?’.

## Notes

### Competing Interest Statement

GJH is a scientific cofounder, and has equity in, Faeth Therapeutics.

### Summary of Updates

New Figure 6 with additional data.

